# Excessive fluid shear stress-mediated Klf4 leads to arteriovenous pathogenesis

**DOI:** 10.1101/2022.07.04.498236

**Authors:** Kuheli Banerjee, Yanzhu Lin, Johannes Gahn, Purnima Gupta, Mariona Graupera, Gergana Dobreva, Martin Schwartz, Roxana Ola

## Abstract

**Background:** Vascular networks form, remodel and mature under the influence of both fluid shear stress (FSS) and soluble factors. For example, FSS synergizes with Bone Morphogenic Protein 9 (BMP9) and BMP10 to promote and maintain vascular stability. Mutation of the BMP receptors ALK1, Endoglin or the downstream effector SMAD4 leads to Hereditary Hemorrhagic Telangiectasia (HHT), characterized by fragile and leaky arterial-venous malformations (AVMs). But how endothelial cells (EC) integrate FSS and BMP signals in normal vascular development and homeostasis, and how mutations give rise to malformations is not well understood.

**Results:** Here we show that loss of *Smad4* in murine ECs increases cells’ sensitivity to flow and the resulting AVMs are characterized by excessive elongation and polarity against the flow. Smad4 deletion also blocks the anti-proliferative response to high FSS, leading to loss of arterial identity. Our data show that loss of cell cycle arrest leads to loss of arterial identity, which is essential in AVM formation upon *Smad4* depletion in ECs. Excessive flow-induced activation of KLF4-PI3K/AKT due to Cyclin dependent Kinase (CDK) activation mediates the aberrant morphological responses to flow triggering AVM formation.

**Conclusions:** Our results show that loss of polarization against the flow is not required for AVM formation upon EC *Smad4* depletion. Instead, increased EC proliferation-mediated loss of arterial identity due to flow-induced PI3K/Akt/Cdks hyperactivation and Klf4 over-expression are the main events associated with AVM formation.

## Introduction

Vascular networks form, remodel and mature under the influence of multiple biomechanical and biochemical signals, but how these are integrated to promote vascular development and maintain adult homeostasis is poorly understood. Fluid shear stress (FSS) from blood flow is a critical variable that determines vascular endothelial cell (EC) number and shape in vascular development and maintenance^1^. ECs also polarize and migrate according to the flow direction; in different systems this may be with or against the flow^2^, but in the developing retina is against the flow, which is proposed to be important in guiding vessel formation^3,4^. One aspect of EC flow responses is the existence of a cell-autonomous shear stress set-point specific to each vessel type. FSS near the set-point promotes EC elongation and alignment parallel to the flow, and stabilizes the vessel whereas flow that is persistently above or below this level triggers vessel remodeling to restore FSS to the appropriate magnitude and contribute to vascular disease^5^.

We previously found that high shear stress within the physiological range synergizes with secreted Bone Morphogenic Protein (BMP) 9 and BMP10 to activate canonical Smad 1/5, which promotes EC quiescence and vascular homeostasis^6^. This pathway contributes to the inhibition of EC proliferation by FSS and to expression of factors that mediate pericyte recruitment, thus, stabilizing the vessels. By contrast, FSS activates the related Smad2/3 pathway only at low FSS magnitude to induce inward arterial remodeling^7^.

Consistent with its role in vascular homeostasis, blocking canonical BMP9/10 signaling in neonatal murine EC results in dilated and leaky arterial-venous malformations (AVMs) in regions of high flow^6,8,9^. These lesions are characteristic of the vascular disorder Hereditary Hemorrhagic Telangiectasia (HHT), an autosomal dominant condition caused by loss-of-function (LOF) heterozygous mutations in the BMP9/10 receptors, Activin Like Kinase 1 (*ALK1*), linked to HHT2, the auxiliary co-receptor-Endoglin (*ENG*), linked to HHT1, and the transcriptional effector-*SMAD4*, linked to Juvenile Polyposis (JP)-HHT^10-12^.

In murine models of HHT, AVMs are characterized by a plethora of dysregulated EC events e.g increased in EC size, misdirected migration, increased proliferation and changes in EC fate ^6,8,9,13-15^. Yet, if one or a complex of interwined cell events, flow dependent or independent drive AVM formation remains largely elusive.

Mechanistically, one important mediator downstream of this flow-BMP9 crosstalk is PI3K/AKT, which is hyperactivated in HHT lesions in mouse models and in human patients^9,14,16^. Pharmacological inhibition of PI3K or depletion of EC *Akt1* rescued AVM formation in HHT murine models^9,14^. Interestingly, PI3K/AKT activated in EC by FSS mediates shear stress-induced EC responses downstream of the mechanosensory junctional receptor complex^17^.

These findings suggest that the canonical BMP9/10-Smad4 signaling plays a crucial role in shear stress regulation of vascular homeostasis. We therefore set out to test the concept that AVMs arise from loss of shear stress-mediated EC quiescence using *Smad4* EC loss of function (LOF) mice as a model of AVM formation.

## Methods

### Animal Experiments

Deletion of endothelial *Smad4* (*Smad4*^*iΔEC*^) *or Klf4* (*Klf4*^*iΔEC*^) was achieved by crossing *Smad4 Fl/Fl* or *Klf4Fl/Fl* with Tx inducible *Cdh5-Cre*^*ERT2*^ mice. To obtain *Smad4;Klf4*^*iΔEC*^ double knockout mice, we crossed *Smad4*^*iΔEC*^ with *Klf4*Fl/Fl mice. Gene deletion was achieved by intra-peritoneal injections of 100µg Tx (Sigma, T5648) into *Smad4*^*iΔEC*^, *Klf4*^*iΔEC*^ and *Smad4*^*iΔEC*^*;Klf4*^*iΔEC*^ at postnatal days (P1-P3). Tx-injected Cre-negative littermates (*Fl/Fl*) were used as controls. The PI3K inhibitor Pictilisib (Selleckchem, S1065, 20 mg/kg/day) and CDK4/6 inhibitor Palbociclib (Selleckchem, S1116, 70 mg/kg/day) were administered intraperitoneally (i.p) at P4 and P5.

Mice were maintained under standard specific pathogen-free conditions, and animal procedures were approved by the animal welfare commission of the Regierungspräsidium Karlsruhe (Karlsruhe, Germany).

### Reagents and antibodies

For immunodetection: anti-VE Cadherin (#2500S, 1:600, Cell Signaling), anti-PECAM (#sc-32732, 1:100 Santa Cruz), Isotectin B4 (IB4, #121412, 10 μg/ml, Life Technologies), anti-GOLPH4 (#ab28049; 1:200, Abcam), anti-GM130 (#610823; 1:600 BD Bioscience), anti-ERG (#92513; 1:200; Abcam), anti-KLF4 (#AF3158, 1:200, R&D systems), anti-phospho S6 (pS6 #5364, 1:200; Cell Signaling), anti-KI67 (eFluor™660, 1:100, ThermoFisher) and anti-SOX17 (#AF1924, 1:200, R&D systems).

For WB: anti-pAKT (#4060, 1:1,000, Cell Signalling), anti-AKT (#4060, 1:1,000, Cell Signalling), anti-SMAD4 (#38454S; 1:1000; Cell Signalling), GAPDH (#5174S 1:5000, Sigma), ACTIN (A1978, 1:1000, Sigma), pRB1 (#8516S, 1:1000, Cell signaling), RB (#9309S, 1:1000, Cell signaling), E2F1 (#3743, 1:1000, Cell signaling), CDK2 (#2546, 1:1000, Cell signaling), CDK6 (#3136, 1:1000, Cell signaling) and CDK4 (#12790, 1:1000, cell signaling).

Appropriate secondary antibodies were fluorescently labelled (Alexa Fluor donkey anti-rabbit, #R37118, Alexa Fluor donkey anti-goat 555, #A-21432, 1:250, Thermo Fisher) or conjugated to horseradish peroxidase for WB (Anti-Rabbit #PI-1000-1, Anti-Goat #PI-9500-1 and Anti-mouse #PI-2000-1 IgG (H+L), 1:5,000, Vector Laboratories).

### mLECs isolation

mLECs were isolated from collagenase I-digested lung tissue using rat anti-mouse CD31 monoclonal antibody-coated Dynabeads (11035, Invitrogen) and directly used for RNA analysis.

### Quantitative real-time PCR

RNAs from HUVECs or mouse lung ECs (mLECs) were purified using RNeasy-kit (74106, Qiagen). The RNA was reverse transcribed High-Capacity cDNA Reverse Transcription Kit (4368813, Thermo Fisher) and quantitative PCR were assayed using PowerUP SYBR Green Master Mix (A25778, Thermo Fisher) with a QuantStudio 3 (Thermo Fisher) according to the manufactures protocol. The following primers were used for mLECs: Klf4 (QT00095431, Qiagen), Smad4 (QT00130585, Qiagen) and Gapdh (Forward: AGGTCGGTGTGAACGGATTTG, Reverse: TGTAGACCATGTAGTTGAGGTCA). Primers used for HUVECs: KLF4 (Forward: CCCACATGAAGCGACTTCCC, Reverse: CAGGTCCAGGAGATCGTTGAA), KLF2 (QT00204729, Qiagen). SMAD4 (QT00013174, Qiagen), PECAM-1 (Forward: AAGTGGAGTCCAGCCGCATATC Reverse: ATGGAGCAGGACAGGTTCAGTC), KDR (QT00069818, Qiagen), CDH5 (QT00013244, Qiagen), GAPDH (Forward: CTGGGCTACACTGAGCACC Reverse: AAGTGGTCGTTGAGGGCAATG), SOX17 (QT00204099, Qiagen), EPRINB2 (forward: TATGCAGAACTGCGATTTCCAA Reverse: TGGGTATAGTACCAGTCCTTGTC).

### Immunostaining

The eyes of P6 pups were fixed in 4% PFA for 17 minutes at room temperature (rt). After several washes with PBS, dissected retinas were incubated with specific antibodies diluted in blocking buffer (1% fetal bovine serum, 3% BSA, 0.5% Triton X-100, 0.01% sodium deoxycholate, 0.02% sodium azide in PBS at pH 7.4) at 4 ° C overnight. The following day, retinas were washed and incubated with IB4 together with the corresponding secondary antibody in PBLEC buffer (1 mM CaCl_2_, 1 mM MgCl_2_, 1 mM MnCl_2_ and 0,25% Triton X-100 in PBS) for 1 hour at rt and mounted in fluorescent mounting medium (RotiMount FluorCare #HP19.1, CarlRoth). High-resolution pictures were acquired using Zeiss LSM800 confocal microscope with Airyscan Detector and the Zeiss ZEN software. Quantification of retinal vasculature was done using Fiji.

### Cell culture, siRNA transfection, overexpression HUVECs and PI3K inhibitor treatment

Human Umbilical Vein Endothelial Cells (HUVECs) were isolated from the umbilical cords of newborn, approved by the local ethics committee (2012-388N-MA, 08/11/2018, Medical Faculty Mannheim, Heidelberg University, Germany). A 3-way valve was inserted into the vein and fixed with a zip tie to wash the vein several times until the effluent buffer was transparent or slightly pink. At that point, the vein was closed with a surgical clamp, filled with a 0.2% Collagenase/Dispase (11097113001, Sigma-Aldrich) solution and incubated at 37°C for 30-45 minutes. Post-incubation the umbilical cord was emptied into 5ml of FCS and centrifuged at 1000rpm for 5 minutes. Then the cells were re-suspended in 10ml of Endothelial Cell Growth Medium MV2 with supplemental mix (C-22022, PromoCell) and 1% Penicillin/Streptomycin (P4333, Sigma-Aldrich), platted on a 10cm dish and incubated at 37°C (with 5% CO_2_, 100% humidity). Cells up to passage 4 were used for experiments. Depletion of *SMAD4, KLF4, PECAM-1, AKT1/2, VEGFR2, CDH5* was achieved by transfecting 25 pmol of siRNA against *SMAD4* (ON-Targetplus Human SMAD4 siRNA Smart Pool, #L-003902-00-0005)), *KLF4* (siGENOME Human KLF4 siRNA; #M-005089-03-0005), PECAM-1 (5’-GGCCCCAAUACACUUCACA-3’), AKT1/2 Stealth siRNA (Thermo Fisher, VHS40082 and VHS41339), VEGFR2 (Dharmacon/Horizon, #L-003148-00-005), CDH5 (Dharmacon/Horizon, #L-003641-00-0005) using Lipofectamine RNAiMax (Invitrogen) in 2% OPTI-MEM. Transfection efficiency was assessed by western blotting and qPCR. Experiments were performed 48-60 hours post transfection and results were compared with siRNA *CTRL* (ON-TARGETplus Non-Targeting Pool D-001810-10-05). Inhibition of PI3K was achieved by using Pictilisib (S1065, Selleckchem) in a concentration of 75nM and inhibition of CDK4/6 by using Palbociclib in concentration of 2 µM. Before experiments cells were starved for 8-10 hours in 2% FCS.

For Generation of stable AKT1 and KLF4 overexpressing cell lines the AKT1 (TRCN0000473539) and KLF4 (TRCN0000492053) overexpression plasmids were obtained from Sigma (Mission TRC3.0, Sigma Aldrich, USA). For lentivirus packing, briefly, HEK293T cells were co-transfected with lentiviral vector and packaging plasmids (pCMV-dR8.91 and pCMV-VSV-G) using X-treme GENE 9 reagent (Sigma). Culture supernatant containing viral particles was collected 36 and 72 h after transfection and concentrated by centrifugation at 3000g for 60 min at 4℃. The pellets were resuspended in 1 mL of PBS and stored at −80℃. For virus infection, HUVECs were transduced with optimal volume of lentiviral virus at 50% confluence in MV2 medium and 8μg/ml Polybrene (Sigma). After 24 h, the medium containing viral particles was replaced with fresh medium and after additional 24h, the infected cells were selected with 2 μg/ml puromycin for 48 hours.

### Exposure of endothelial cells to increased shear stress

HUVECs transfected with siRNAs or OE-HUVECs were plated in a six-well plate and on an orbital shaker (Rotamax120, Heidolph Instruments) at 50, 150 or 250 rpm to generate laminar sheer stress of 1, 5 or 12 DYNES/cm^2^ respectively. Results were confirmed in a µ-Slide VI^0.4^ (Ibidi, 80601) using a pump system (Ibidi, 10902).

### Western blotting

HUVECs were washed with PBS and lysed with Laemmli buffer (1610740, Biorad). Samples were separated on 10% SDS-PAGE gels and transferred on 0.2µm nitrocellulose membranes (10600004, GE Healthcare). Western blots were developed with the Clarity Western ECL Substrate (1705061, Biorad) on a Luminescent image Analyzer, Fusion FX (Vilber). Bands’ intensity were quantified using ImageJ.

### Proliferation Assay

Proliferation analysis was performed using Click-iT EdU Alexa Fluor 488 Imaging kit (Life Technologies). P6 pups were injected with 200 μg of EdU (5 mg/mL) and sacrificed 4 hours later. EdU staining was done according the manufacturer’s protocol.

### RNA-Sequencing

For RNA-Seq of HUVECs the RNA was isolated with the Quick-RNA Miniprep Kit (#R1054, Zymo Research), 60 hours after transfection of Control or Smad4 siRNA. The RNA 6000 Nano Kit (#5067-1511, Agilent) was used to assess the RNA integrity on a Bioanalyzer 2100 (Agilent). Both sequencing and library preparation were performed on the BGISEQ-500 platform.

### RNA-Sequencing data analysis

Quality of RNA seq reads was assessed with the MultiQC tool (v1.13) and trimmed of adapters using Trimmomatic (v0.39). Reads were mapped by STAR (v2.7.10a) with the following settings: -alignIntronMin 20 and -alignIntronMax 500000 to the hg38 reference genome. Tag directories were created with makeTagDirectory and reads were counted by the analyzeRepeats.pl function (rna hg38 -strad both -count exons -noadj) both from HOMER (v4.7.2). Differential expression was quantified and normalized using DESeq2. Rpkm.default from EdgeR was used to determine average reads per millions mapped (RPKM). Heatmaps were created by using heatmapper.ca from the RPKM values and represent the row-based Z-scores.

### Statistical analysis

All data are shown as mean ± standard error of the mean (SEM). Samples with equal variances were tested using Mann–Whitney U test or two-tailed Student’s t-test between groups. P value <0.05 was considered to be statistically significant. Statistical analyses were performed for all quantitative data using Prism 9.0 (Graph Pad).

## Results

### Smad4 signaling maintains the shear stress set-point-mediated EC responses

Impaired responses of EC to FSS, including migration direction and changes in EC size, have been proposed to mediate HHT lesions, mainly in models of HHT1 and HHT2, ie., mutations in *ENG* and *ALK1*^8,15,18^. To explore flow-mediated EC events in JP-HHT where *SMAD4* is mutated, we depleted primary human umbilical cord ECs (HUVECs) of *SMAD4* using small interfering RNA (siRNA) versus *CTRL* siRNA (confirmed in **Figure 1G**). Cells were subject to laminar shear stress at 1 or 12 DYNES/cm^2^ for 24 and 48 hours (**Figure 1A-I**). *SMAD4*-depleted HUVECs were more elongated without flow (0 hours) (**Figure 1A,D**). At 24 hours under 12 DYNES/cm^2^, cells elongated further and aligned better in the direction of flow compared to controls (**Figure 1B,E**; quantified in **1H,I)**. Under 1 DYNE/cm^2^ stress, *CTRL* HUVECs failed to elongate or align even at 48 hours, whereas *SMAD4*-depleted HUVECs showed distinct elongation and alignment over this time (**Figure 1C,F,I**). Thus, flow-induced EC elongation and alignment are both faster and more sensitive after *SMAD4* knock down (*SMAD4*KD).

**Figure 1.**
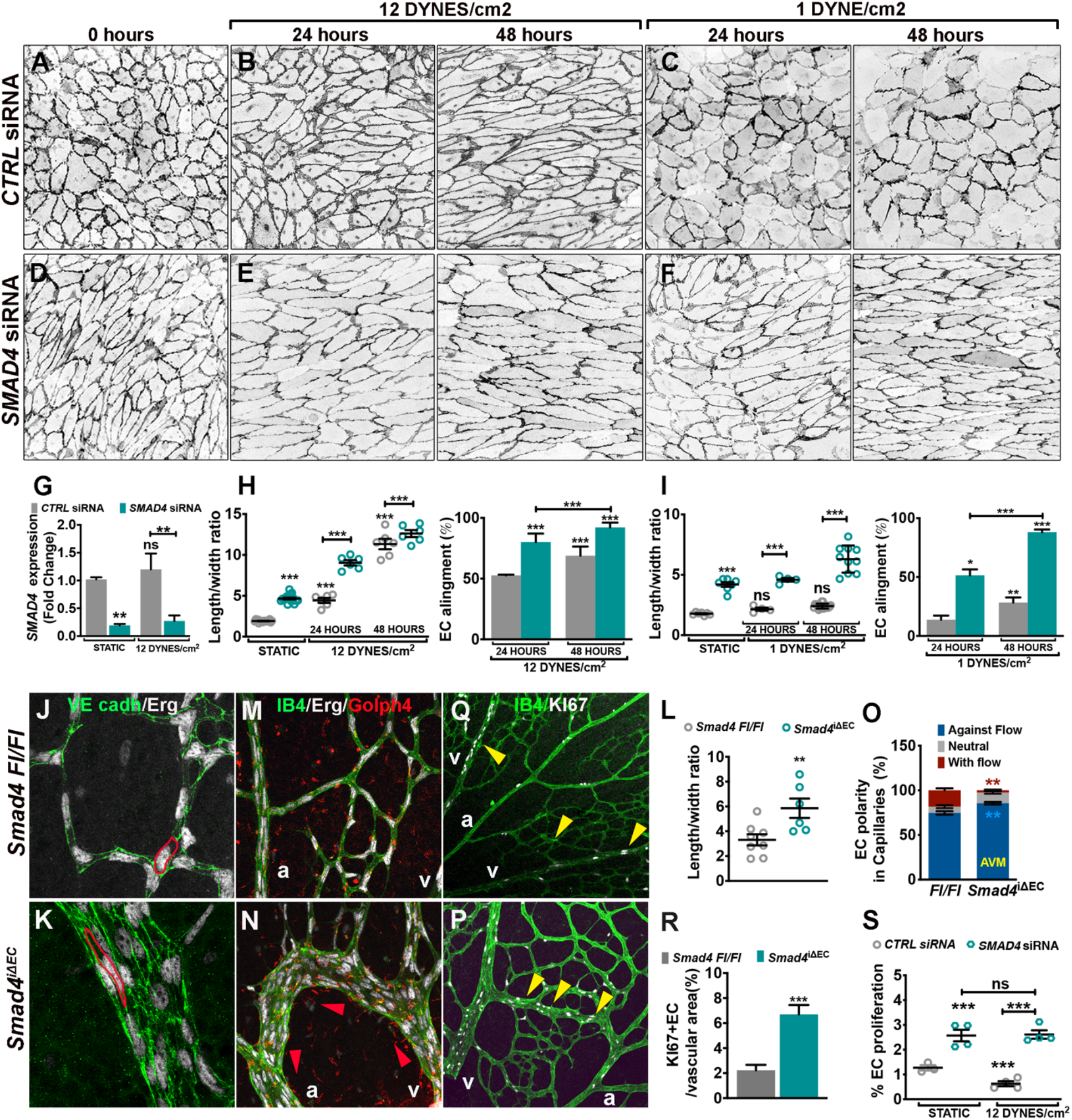
SMAD4 signaling maintains the FSS set-point to restrict flow mediated EC responses. (**A**-**F**) Negative images of VE Cadherin staining of HUVECs transfected with *CTRL* (**A-C**) and *SMAD4* (**D**-**F**) *siRNAs* grown in static (**A,D**) or increasing flow magnitudes: 12 DYNES/cm^2^ (**B,E**) or 1 DYNE/cm^2^ (**C,F**) for 24 and 48 hours, respectively. The direction of the flow is right to left. (**G**) *SMAD4* qPCR expression (fold change) in *CTRL* versus *SMAD4* siRNAs HUVECs. (**H,I**) Quantification of length/width ratio and of EC alignment parallel to flow direction (%) in *CTRL* and *SMAD4* siRNAs HUVECs grown in static versus subjected to 12 DYNES/cm^2^ (**H**) and 1 DYNE/cm^2^ (**I**). (**J,K**) Labeling of postnatal day 6 (P6) Tx induced *Smad4 Fl/Fl* (**J)** and *Smad4*^i∆EC^ (**K**) retinas for Erg (labelling the EC nuclei-white), and VE-Cadherin (green). (**L**) Quantification of length/width ratio in capillaries versus AVMs in *Smad4 Fl/Fl* and *Smad4*^i∆EC^ retinas. (**M,N**) Labeling of P6 Tx induced *Smad4 Fl/Fl* and *Smad4*^i∆EC^ retinas for Erg (labelling the EC nuclei-white), Golph4 (labeling the Golgi-red) and Isolectin (IB4) (labeling vessels-green). (**O**) Quantification of EC polarization against versus towards versus non-oriented (neutral) in the capillaries of *Smad4 Fl/Fl* and *Smad4*^i∆EC^ retinas. (**Q,P**) Labeling for KI67 (white) and IB4 (green) of the vascular plexus from P6 *Smad4 Fl/Fl* and *Smad4*^i∆EC^ retinas. (**R**) Quantification of the number of KI67 positive nuclei per vascular area in the vascular plexus (%). (**S**) EC proliferation in response to 12 dynes/cm^2^ for 24 hours of *CTRL* and *SMAD4* siRNA HUVECs. Cell proliferation was measured by incorporation of EdU for 4 hours. Scale Bars: 100μm in **A**-**F** and **M,N 200**μm in **J,K** and 50μm in **Q,P**. Red arrows in **N** point to the direction of polarization against the flow. Yellow arrows in **Q,P** point to KI67+ ECs in the vascular plexus. **a**: artery, **v**: vein. Error bars: s.e.m., \**P*<0.05,\*\**P*<0.01,\*\*\**P*<0.001, ns- non-significant student T test.

To test these observations *in vivo*, we measured the length/width ratio of individual EC labelled for VE cadherin and Erg within the capillaries in *Cdh5-Cre* negative (*Smad4 Fl/Fl* - control) versus AVMs in tamoxifen-inducible *Smad4* EC-specific deficient postnatal day 6 (P6) (*Smad4*^*iΔEC*^ or *Smad4*ECko) retinas. These measurements confirmed increased morphological EC responses to flow upon *Smad4* depletion (**Figure 1J,K**; quantified in **1L**). Together, these results imply that SMAD4 signaling restricts shear stress-mediated EC shape responses to flow.

ECs in the postnatal retina polarize and migrate against the flow direction (axial polarity), from the veins towards the arteries, with the degree of polarization correlating with shear stress magnitude^3^. Disrupted polarization and impaired movement of ECs against the direction of flow has been proposed to mediate AVM formation in *Eng* and *Alk1* mutants^8,15^. We therefore analyzed polarity in capillaries versus AVMs in *Smad4 Fl/Fl* and *Smad4*^*iΔEC*^ P6 retinas by staining for Golph4 to label the Golgi apparatus, Erg for the EC nuclei and Isolectin B4 (IB4) to visualize the endothelium (**Figure 1M,N**). The relative position of the Golgi and nuclei were then quantified (**Figure 1O**). EC polarization against the predicted flow direction was significantly increased in AVMs in *Smad4*^*iΔEC*^ retinas (**Figure 1N,O**). Thus, multiple EC morphological responses to shear stress are increased after *SMAD4* KD *in vitro* or *Smad4*ECko *in vivo*.

It is well established that physiologically high FSS inhibits EC proliferation^19^. As expected^9^, labelling of *Smad4 Fl/Fl* and *Smad4*^*iΔEC*^ retinas for the mitotic marker KI67 and the total EC marker IB4 revealed increased EC proliferation within AVMs (**Figure 1Q,P**; quantified in **1R**). *In vitro*, EdU labeling to identify ECs in S phase showed that *SMAD4* depletion increased baseline cell cycle progression and completely blocked the inhibition by high shear stress (**Figure 1S**). Thus, Smad4 is required for flow-mediated repression of EC proliferation.

Taken together, these results show that Smad4 resembles Alk1 and Eng in that it is also required for flow-mediated EC proliferation but is opposite in that it suppresses rather than enhances morphological responses to flow.

### KLF4 mediates flow-induced hyper-responsiveness of *SMAD4* depleted HUVECs

To identify mediators of increased responsiveness of *SMAD4*ECko to FSS we performed RNA sequencing in *CTRL* versus *SMAD4* siRNAs HUVECs grown in static or subjected to 2 hours 12 DYNES/cm^2^ and focused on shear stress responsive genes. Interestingly, among other flow regulated genes, *SMAD4* depletion enhanced FSS-induced Krüppel-like transcription factor (KLF) 2 and KLF4 induction (**Figure 2A**). As the two mechanosensors show dose-dependent induction by FSS^20^, and to validate our transcriptomic results, we perfomed RT-PCR for *KLF2* and *KLF4* in HUVECs subjected to increasing magnitudes of laminar FSS (1-5-12 DYNES/cm^2^) and depleted for *SMAD4*. Interestingly, loss of *SMAD4* moderately augmented flow-induced KLF2 and KLF4 with increasing flow magnitude (**Figure 2B**).

**Figure 2.**
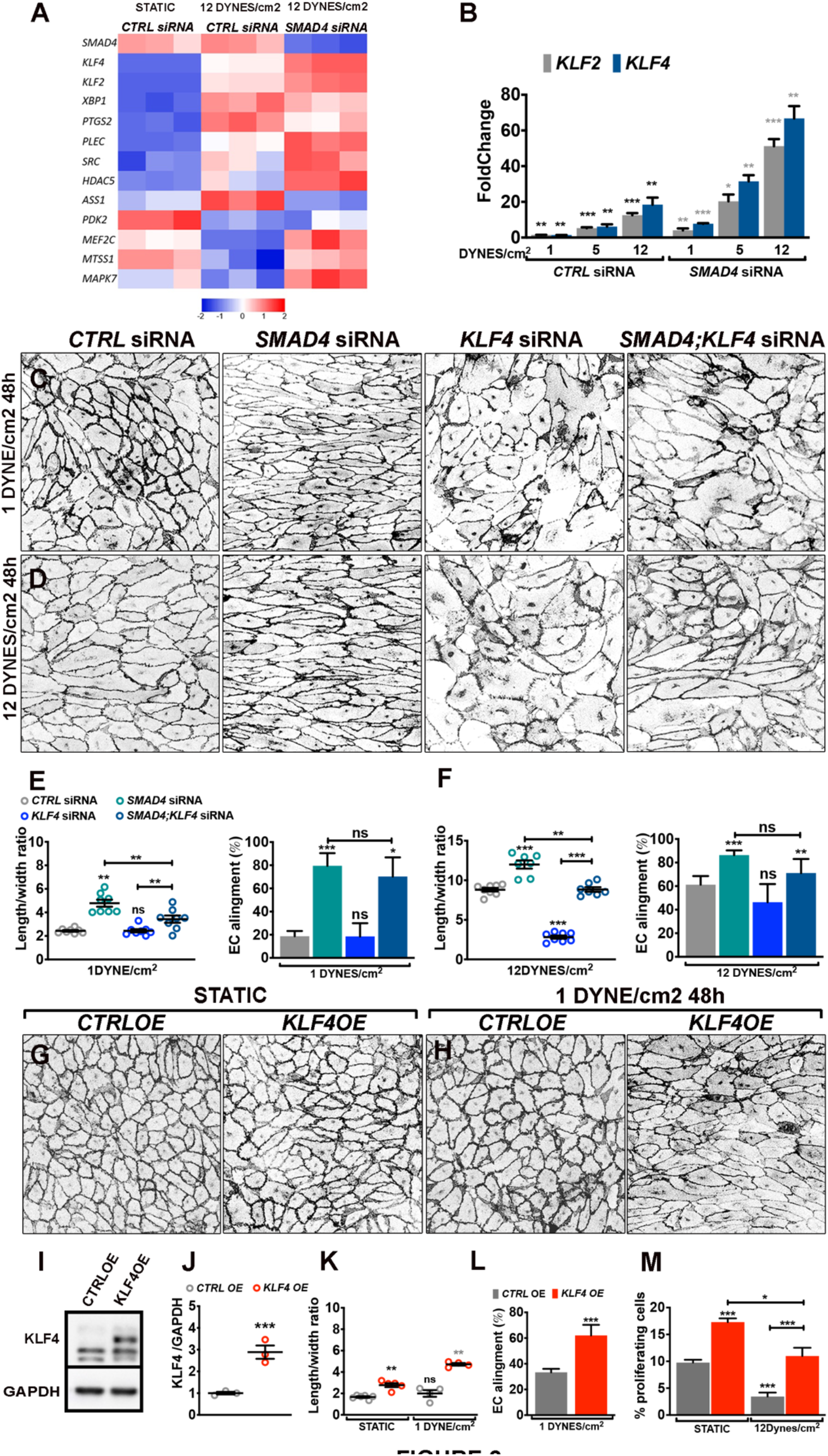
KLF4 mediates the hyper-responsiveness of *SMAD4* depleted cells upon flow. **(A)** Heatmap for the expression of FSS responsive genes in *CTRL* versus *SMAD4* siRNA HUVECs subjected to 12DYNES/cm^2^; n = 3 samples in each group. Color key shows log2 change after SMAD4 depletion. **(B)** *KLF*2 and *KLF4* mRNA expression by qPCR in HUVECs transfected with *CTRL* and *SMAD4* siRNAs grown in static or subjected to increasing magnitudes of shear stress: 1-5-12 DYNES/cm^2^ for 2 hours. (**C,D**) Negative images of VE Cadherin staining of HUVECs transfected with *CTRL, SMAD4, KLF4* and *SMAD4;KLF4* siRNAs subjected to 1 DYNE/cm^2^ (**C**) and 12 DYNES/cm^2^ (**D**) for 48 hours. (**E,F**) Quantification of the length/width ratio and of EC alignment parallel to the flow direction (%) of HUVECs transfected with *CTRL, SMAD4, KLF4* and *SMAD4;KLF4siRNAs* subjected to 1 DYNE/cm^2^ (**C**) and 12 DYNES/cm^2^ (**D**). (**G,H**) Negative images of VE Cadherin staining of HUVECs transfected with *CTRL*OE and *KLF4*OE constructs grown in static (**G**) or subjected to 1DYNE/cm^2^ for 48 hours (**H**). (**I**) WB analysis of HUVECs transfected with an empty lentiviral construct (*CTRL*OE) and an overexpression lentivirus for *KLF4* (*KLF4*OE). (**J**) Quantification of KLF4 protein expression levels normalized to GAPDH. (**K,L**) Quantification of the length/width ratio (**K**) and of EC alignment (%) parallel to the flow (**L**) of *CTRL*OE and *KLF4*OE HUVECs grown in static versus 1 DYNE/cm^2^ conditions for 48 hours. Scale Bars: 100μm in **A,B** and **G,H**. Error bars: s.e.m., n.s- non-significant, \**P*<0.05, \*\**P*<0.01, \*\*\**P*<0.001, student T test.

As KLF4 showed the highest induction to FSS upon *SMAD4* depletion in both transcriptomics and RT-PCR data, we therefore considered the role of KLF4 in the altered behaviours of *SMAD4*KD cells. HUVECs were depleted for either *SMAD4* or *KLF4* or both and subject to low versus high FSS for 48 hours. *KLF4*KD failed to elongate but aligned under flow and *KLF4* inactivation reversed the highly elongated morphology of *SMAD4* depleted HUVECs under all conditions with no effect on cell alignment (**Figure 2C,D**; quantified **in 2E,F**).

Conversely, we overexpressed *KLF4* using a lentiviral vector in HUVECs (*KLF4*OE; confirmed in **Figure 2I,J**) and subjected cells to low FSS for 48 hours. *KLF4*OE increased cell elongation under static conditions (**Figure 2G**; quantified in **2K**), which was further enhanced by low FSS (**Figure 2H,K**). *KLF4*OE cells also aligned better in the flow direction (**Figure 2H**; quantified in **2L**). Measurement of EdU incorporation showed that *KLF4*OE increased cell cycle progression both at baseline and under high FSS (**Figure 2M**). *KLF4*OE thus induces many of the key effects of *SMAD4*KD. Together, these results show that KLF4 contributes to the morphological effects mediated by SMAD4 LOF.

### High flow-induced KLF4 is a key determinant in AVM formation

To gain insights into Smad4 regulation of flow-induced Klf4 *in vivo*, we labelled *Smad4* Fl/Fl and *Smad4*^*iΔEC*^ retinas for Klf4 and for IB4 (**Figure 3A-D’**). Within the retinal developing plexus, the shear stress levels are the highest in the vascular plexus close to the optic nerve, and gradually decrease toward the sprouting front^21,22^. In *Smad4* Fl/Fl retinas, Klf4 expression was minimal in the low shear vascular front and capillary ECs (**Figure 3A**), moderate in higher flow large veins, increased further in larger arteries, and reached the highest intensity at the first retinal branch points where the wall shear stress is maximal (**arrows in Figure 3B,B’**). Interestingly, this specific region corresponds to the most frequent site of AVM formation^9^. In *Smad4*^*iΔEC*^ retinas, *Klf4* expression was highly upregulated in AVMs at the highest intensity relative to the feeding artery and vein (red arrows in **Figure 3D** and **Figure 3D’**, quantified in **3E**). The arteries and veins upstream of AVMs (yellow arrows in **Figure 3D,D’**) or vessels not engaged in AVMs (white arrows in **Figure 3D,D’**) showed lower Klf4 intensity. This finding is consistent with recent findings showing lower flow outside of AVMs in embryos with decreased Cx37^23^. It also suggests that FSS-induced *Klf4* expression is partially restrained by Smad4, thus confirming our *in vitro* data. These results emphasize that high Klf4 within AVM ECs is likely a consequence of both increased sensitivity and high flow.

**Figure 3.**
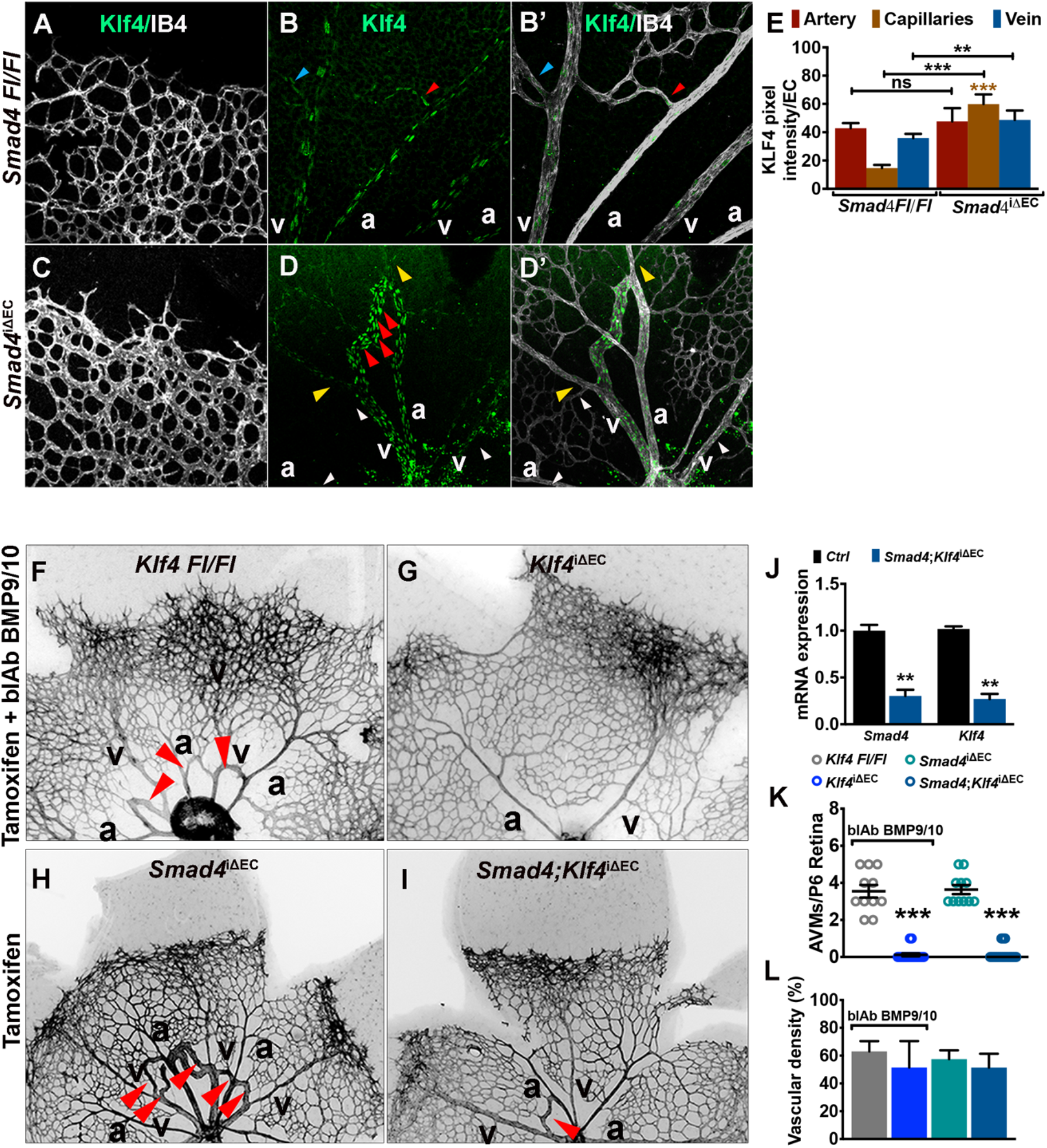
High Flow-induced KLF4 is a key determinant in AVM formation. (**A**-**D’**) Labeling of Tx induced P6 *Smad4* Fl/Fl and *Smad4*^i∆EC^ retinas for Klf4 (green) and IB4 (white) in the sprouting front (**A,C**) versus vascular plexus (**B,B’** and **D,D’**). Small red/blue arrowheads in **B,B’** indicate the first branch points in artery/vein. Red arrowheads in **D,D’** indicate increased KLF4 intensity within the AVM. Small yellow/white arrowheads indicate vessels upstream of AVMs or vessels not engaged in AVMs expressing very little KLF4. (**E**) Quantification of KLF4 pixel intensity/EC in arteries, capillaries and veins in Fl/Fl and *Smad4*^i∆EC^ retinas. (**F,G,H,I**) Staining of P6 retinas with IB4 (negative images) of *Klf4 Fl/Fl* (**F**) and *Klf4*^i∆EC^ (**G**) treated with blocking antibodies for BMP9/10 (BMP9/10blAb) and of *Smad4*^i∆EC^ (**H**) and double knockout mice:*Smad4;Klf4*^i∆EC^(**I**). Red arrows in **F, H** and **I** mark AVMs. **(K**) *Smad4* and *Klf4* mRNA expression by qPCR in purified mouse lung endothelial cells (mLECs) isolated from P6 Tx injected mice. (**L**) Quantification of P6 retinal AVMs’ number. (**M**) Quantification of vascular density at the retinal front (%). Scale Bars: 100μm in **A-D**’ and 20μm in **G,H,I,J**. Error bars: \*\**P*<0.01, \*\*\**P*<0.001, ns- non-significant, student T test.**a**: artery, **v**: vein.

To test Klf4 function *in vivo*, we generated two genetic models. First, we examined EC specific Tx-inducible *Klf4* LOF neonates (*Klf4*^i∆EC^) where AVMs were induced by administration of blocking antibodies for BMP9/10 (**Figure 3F,G**). Second, we created EC specific Tx-inducible double ko mice, *Smad4;Klf4*^i∆EC^ (**Figure 3H,I**). Tx was injected at P1-P3 and retinas were analysed at P6. Efficient *Smad4* and *Klf4* gene deletion was validated by qPCR from P6 mouse lung endothelial cells (mLECs; **Figure 3J**). Blockade of BMP9/10 in control *Klf4 Fl/Fl* retinas led to formation of AVMs (average of 3.6-4 AVMs per retina) and an increase in vascular front density similar to *Smad4*ECko (**Figure 3F,H**; quantified in **3K,L**). *Klf4* inactivation rescued AVM formation in both models but not the excessive sprouting at the vascular front (**Figure 3G,I**; quantified in **3K,L**). Thus, flow-induced Klf4 is a key determinant in AVM pathogenesis and the first molecular marker identified to-date to distinguish flow-dependent AVM formation from flow-independent excessive sprouting.

### KLF4 mediates the shear stress-induced aberrant EC events within AVMs

To address the role of Klf4 in EC axial polarity, *Fl/Fl, Smad4*^*iΔEC*^, *Klf4*^*iΔEC*^ and *Smad4*;*Klf4*^iΔEC^ retinas were examined for Golgi orientation. Compared to *Fl/Fl* mice, *Klf4* deficient capillaries showed reduced polarization against the flow direction; in *Smad4*^*iΔEC*^ retinas, additional *Klf4* inactivation blunted the increased axial polarity (**Figure 4A,B**; quantified in **Figure 4C**). *Klf4*ko ECs were less elongated than in *Fl/Fl*, and *Klf4* inactivation rescued the excessive elongation of *Smad4*^*iΔEC*^ ECs, confirming our *in vitro* findings (**Figure 4D**). To assay Klf4-mediated EC cell cycle progression, we injected EdU into *Fl/Fl*, S*mad4*^iΔEC^ and S*mad4*;*Klf4*^iΔEC^ P6 pups, 4 hours before collecting tissue and labeling for EdU and Erg (**Figure 4E**). As previously observed, EdU+/Erg+ double positive ECs increased markedly in *Smad4*^*iΔEC*^ retinas, exclusively in AVMs. *Klf4* inactivation significantly decreased the number of EdU+/Erg+ in the vascular plexus of *Smad4*^iΔEC^ retinas to levels comparable to *Fl/Fl* retinas (**Figure 4E**; quantified in **4F)**. Elevated Klf4 thus contributes to increased morphological responses and excessive proliferation in *Smad4*ECko AVMs.

**Figure 4.**
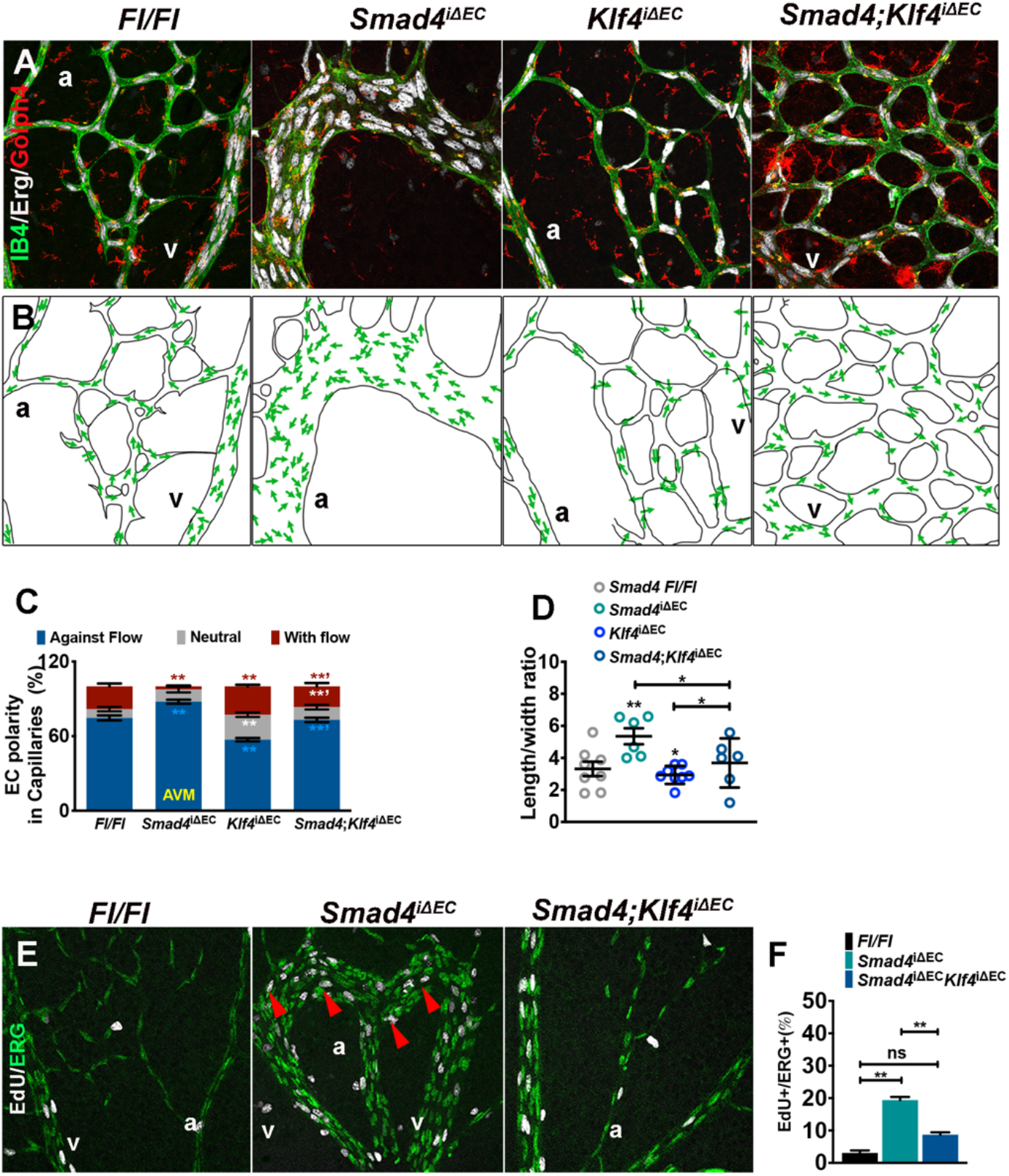
KLF4 mediates the shear stress-induced aberrant EC events within AVMs. (**A**) Confocal images of P6 Tx induced *Fl/Fl, Smad4*^i∆EC^, *Klf4*^i∆EC^ *and Smad4;Klf4*^i∆EC^ retinal plexus labeled for Erg (white), Golph4 (Golgi-red) and IB4 (green). (**B**) Panels illustrating EC polarization based on position of Golgi versus nucleus in the direction of migration (green arrows). (**C**) Quantification of EC polarization: against the direction of flow, towards the direction of flow and neutral in capillaries from P6 Tx induced retinas from the indicated genotypes. (**D**) Quantification of length/width ratio in capillary ECs in the indicated genotypes. (**E**) Labeling for EdU (white) and Erg (green) in vascular plexus of retinas from P6 *Fl/Fl, Smad4*^i∆EC^ and *Smad4;Klf4*^i∆EC^ mice. Red arrowheads indicate the EdU+ /Erg+ cells in the AVMs. (**F**) Quantification of the number of Erg/EdU double + EC nuclei in the vascular plexus (%). Scale Bars: 100μm in **A,B,E**. Error bars: n.s-non-significant, \**P*<0.05, \*\**P*<0.01, \*\*\**P*<0.01, student T test. **a**: artery, **v**: vein.

### Excessive flow-mediated PI3K/AKT activation regulates flow-mediated EC responses

We previously identified increased PI3K/AKT activity upon inactivation of BMP9/10-Alk1-Smad4 in ECs, further augmented by high FSS^9,14^ and PI3K downstream of flow mediates EC responses^17^. To further understand if increased responsiveness of *SMAD4* deficient cells to FSS is due to PI3K/AKT pathway activation, we subjected *CTRL* versus *SMAD4* siRNAs HUVECs to increasing magnitudes of shear stress (1-5-12 DYNES/cm^2^). Interstingly, *SMAD4* deletion significantly increased AKT phosphorylation at serine 473, a marker of activation, under static condition and increasing flow magnitudes had an additive effect (**Figure 5A**; quantified in **5B**).

**Figure 5.**
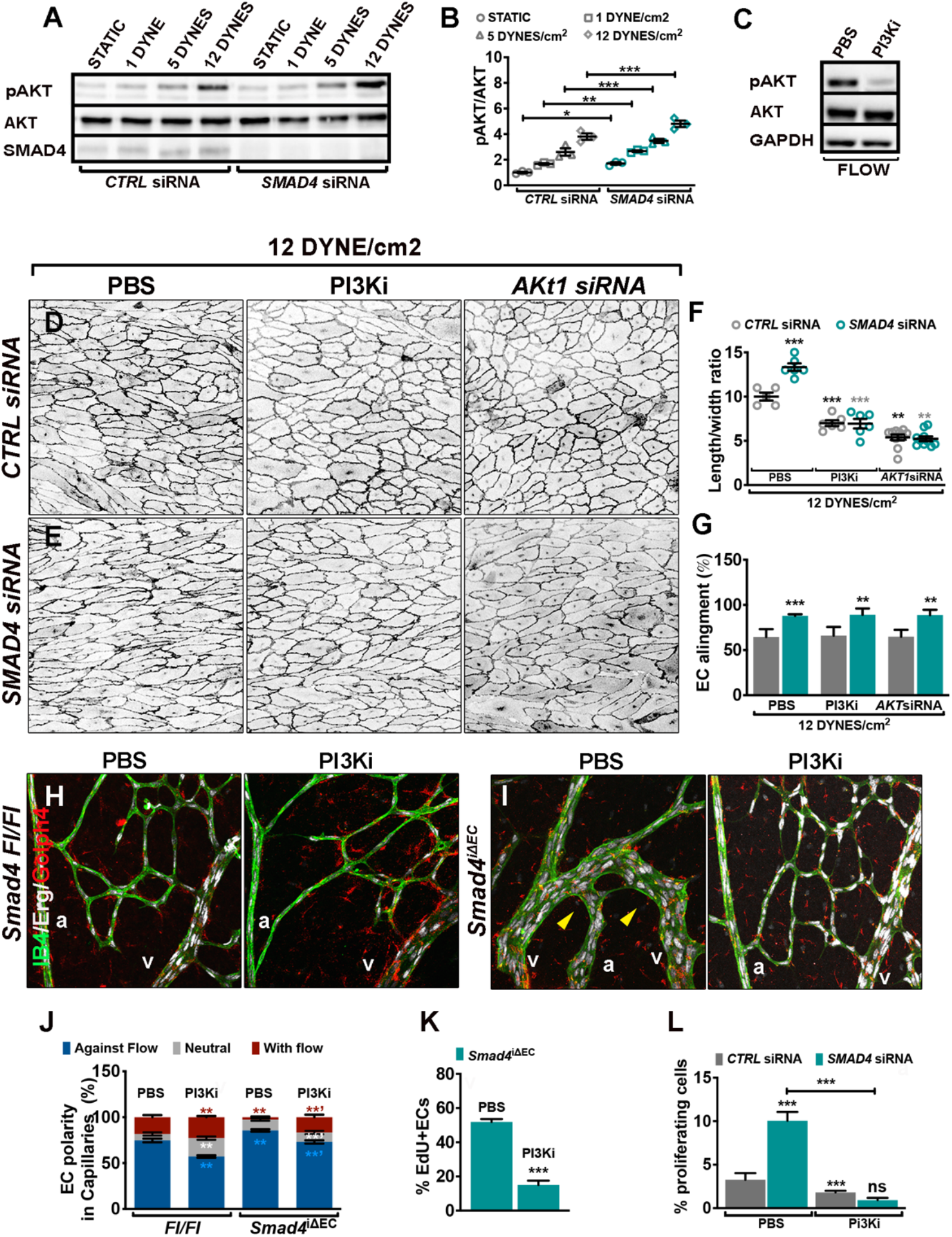
Smad4-induced PI3K/AKT activation regulates flow-mediated EC responses. (**A**) Western Blot (WB) analysis of HUVECs transfected with *CTRL* or *SMAD4* siRNA grown in static or subject to increasing magnitudes of shear stress: 1-5-12 DYNES/cm^2^ for 4 hours. (**B**) Quantifications of pAKT levels normalized to total AKT. (**C**) WB analysis of HUVECs subject to 5 DYNES/cm^2^ treated with PBS or Pictilisib (PI3Ki-75nM) for 4 hours. (**D,E**) Negative images of VE-Cadherin staining of HUVECs transfected with *CTRL* and *SMAD4* siRNAs subjected to 12 DYNE/cm^2^ and treated with PI3K inhibitor or with *AKT1* siRNA for 48 hours. (**F,G**) Quantification of the length/width ratio (**F**) and of EC alignment parallel to flow direction (%) (**G**). (**H,I**) Confocal images of retinas from P6 *Fl/Fl* (**H)** and *Smad4*^i∆EC^ (**I**) from pups treated with PBS or PI3K inhibitor labeled for Erg (white), Golph4 (Golgi-red) and IB4 (green). Yellow arrows in **I** mark the AVMs. (**J**) Quantification of EC polarization: against the direction of flow, towards the direction of flow and non-oriented (neutral) in capillaries and AVMs from P6 retinas of *Smad4Fl/Fl* and *Smad4*^iΔEC^ pups treated with PBS or PI3Ki. (**K**) Quantification of EdU+/Erg+ ECs in the vascular plexus of *Smad4*^iΔEC^ retinas in PBS versus PI3Ki (Pictilisib) treated pups (**L**) EC proliferation (incorporation of EdU) in response to 24 hours 12 DYNES/cm^2^ of *CTRL* and *SMAD4* siRNAs HUVECs treated with PBS versus PI3Ki treatment. Scale Bars: 100μm in **D,E,H, I**. Error bars: n.s-non-significant, \**P*<0.05, \*\**P*<0.01, \*\*\**P*<0.01, student T test. **a**: artery, **v**: vein.

To assess the role of activated AKT in the amplified morphological response to FSS, we inhibited PI3K/AKT signaling for 48 hours using a specific PI3K inhibitor-Pictilisib (confirmed by Western Blot (WB)) (**Figure 5C**), or by transfection with *AKT1* siRNA. Inhibition of PI3K-AKT by either method significantly rescued the length/width ratio in *SMAD4* depleted HUVECs without affecting EC alignment (**Figure 5D,E**; quantified in **5F,G**).

To test *in vivo*, we treated *Smad4 Fl/Fl* and *Smad4*^*iΔEC*^ pups with Pictilisib and examined retinas labeled with IB4, Erg and Golph4. Here, we found that PI3K inhibition blunted the axial polarity in both *Smad4 Fl/Fl* and *Smad4*^*iΔEC*^ retinas (**Figure 5 H,I**; quantified in **5J**). EdU labeling in these mice showed that inhibition of AKT rescued the excessive EC proliferation in *Smad4*^*iΔEC*^ vascular plexus ECs (**Figure 5K**). *In vitro*, inhibition of PI3K also reversed the excess cell cycle progression after *SMAD4KD in* ECs under high FSS (**Figure 5L**). Thus, endothelial SMAD4 functions to restrain flow-induced KLF4 and PI3K/AKT and downstream responses including elongation, axial polarity and proliferation but not the EC alingment.

### Flow-induced KLF4 acts upstream of mechanosensory complex-PI3K/AKT pathway

To untangle the relationships between KLF4 and PI3K in the context of SMAD4 depletion and flow, we subjected HUVECs to flow in the presence of Pictilisib. RT-PCR results show no effect of Pictilisib on flow-induced *KLF4* (**Figure 6A**). We also considered the role of the junctional mechanosensory receptor complex that mediates flow responses including PI3K activation^24,25^. Depletion of each of the components of the mechanosensory receptor complex had no effect on the flow-upregulation of *KLF4* expression (**Figure 6B**). Thus, flow-induced *KLF4* expression does not require PI3K or the mechanosensory junctional receptor complex.

**Figure 6.**
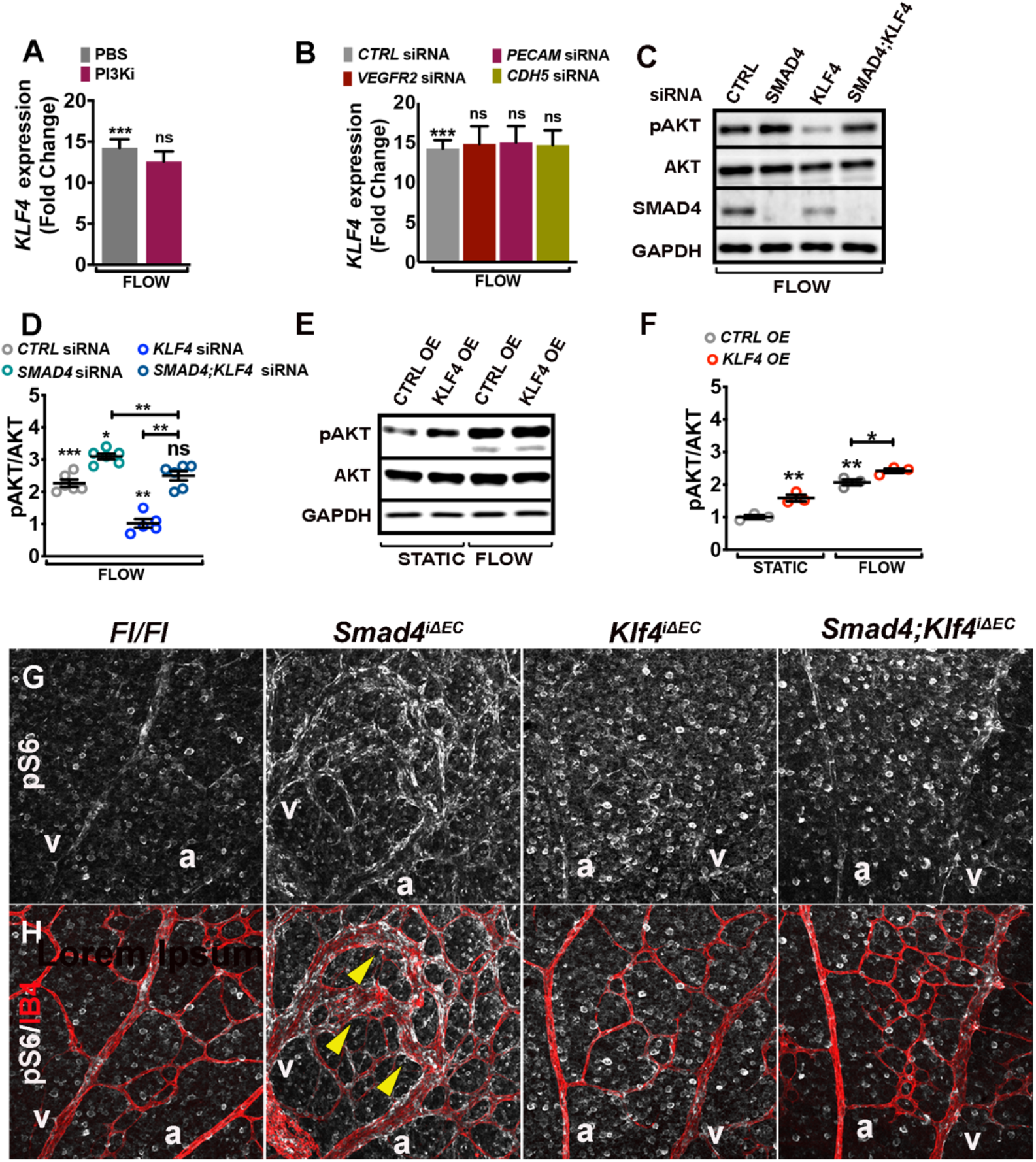
Flow-induced KLF4 acts upstream of mechanosensory complex-PI3K/AKT pathway activation. (**A,B**) *KLF4* mRNA expression by qPCR in HUVECs subjected to 5 DYNES/cm^2^ and treated with PBS versus Pictilisib (PI3Ki-75nM) (**A**) and in *CTRL, VEGFR2, PECAM* and *CDH5* siRNAs HUVECs (**B**) subjected to 5 DYNES/cm^2^ for 2 hours. (**C**) WB analysis for pAKT, total AKT, SMAD4 and GAPDH of *CTRL, SMAD4, KLF4* and *SMAD4;KLF4* siRNAs HUVECs subjected to 5 DYNES/cm^2^ for 4 hours. (**D**) Quantification of pAKT levels normalized to total AKT. (**E**) WB analysis of HUVECs transfected with an empty lentiviral construct (*CTRL-OE*) and an overexpression lentivirus for *KLF4* (*KLF4-OE*). (**F**) Quantification of pAKT levels normalized to total AKT. (**G,H**) Anti-pS6 (white) alone (**G**) and double labeling for pS6 (white) and IB4 (red) staining (**H**) of retinal flat mounts from Tx induced P6 *Fl/Fl, Smad4*^iΔEC^, *Klf4*^iΔEC^ and *Smad4;Klf4*^iΔEC^. Scale Bars: 100μm in **G,H**. Error bars: s.e.m., ns: non-significant, \**P*<0.05, \*\**P*<0.01, \*\*\**P*<0.001, student T test.

As both, FSS-induced *Klf4* expression and AKT activation are partially restrained by Smad4 in a similar manner, we then tested effects of KLF4 on AKT activation, with flow and *SMAD4*KD. *KLF4* inactivation blunted the increase in AKT activity under flow and rescued AKT hyperactivation in *SMAD4* depleted HUVECs (**Figure 6C**; quantified in **6D**). To further test if flow-induced KLF4 is upstream of PI3K we examined the *KLF4*OE HUVECs. KLF4 upregulation was sufficient to activate AKT, with or without FSS (**Figure 6E**; quantified in **6F**). To test these findings *in vivo*, we labelled retinas for phosphorylated S6 ribosomal protein (pS6), a downstream target of AKT activation, and with IB4 to label ECs. ECs in *Smad4*^iΔEC^ retinas showed high pS6 as expected, which was largely rescued by *Klf4* deficiency (**Figure 6G,H**). Klf4 is thus upstream of PI3K to control flow-mediated EC events after *Smad4*ECko.

### Increased EC proliferation-mediated loss of arterial identity is the main driver of AVMs

Current models propose that decreased polarization and migration of ECs against the direction of flow is critical in AVM formation upon *Eng* and *Alk1ECko*^8,15^. The data herein shows that AVM formation upon *Smad4* depletion involves, if anything, increased polarity. These findings prompted us to investigate if instead, increased EC proliferation is the main event triggering AVMs.

Cell cycle distribution in fluorescence activated cell sorted (FACS) ECs from P6 *Smad4 Fl/Fl* and *Smad4*^*iΔEC*^ retinas revealed an increase in actively cycling ECs in S/G2/M together with a decrease in ECs in G1, confirming increased EC proliferation upon *Smad4*ECko (**Figure 7A**).

**Figure 7.**
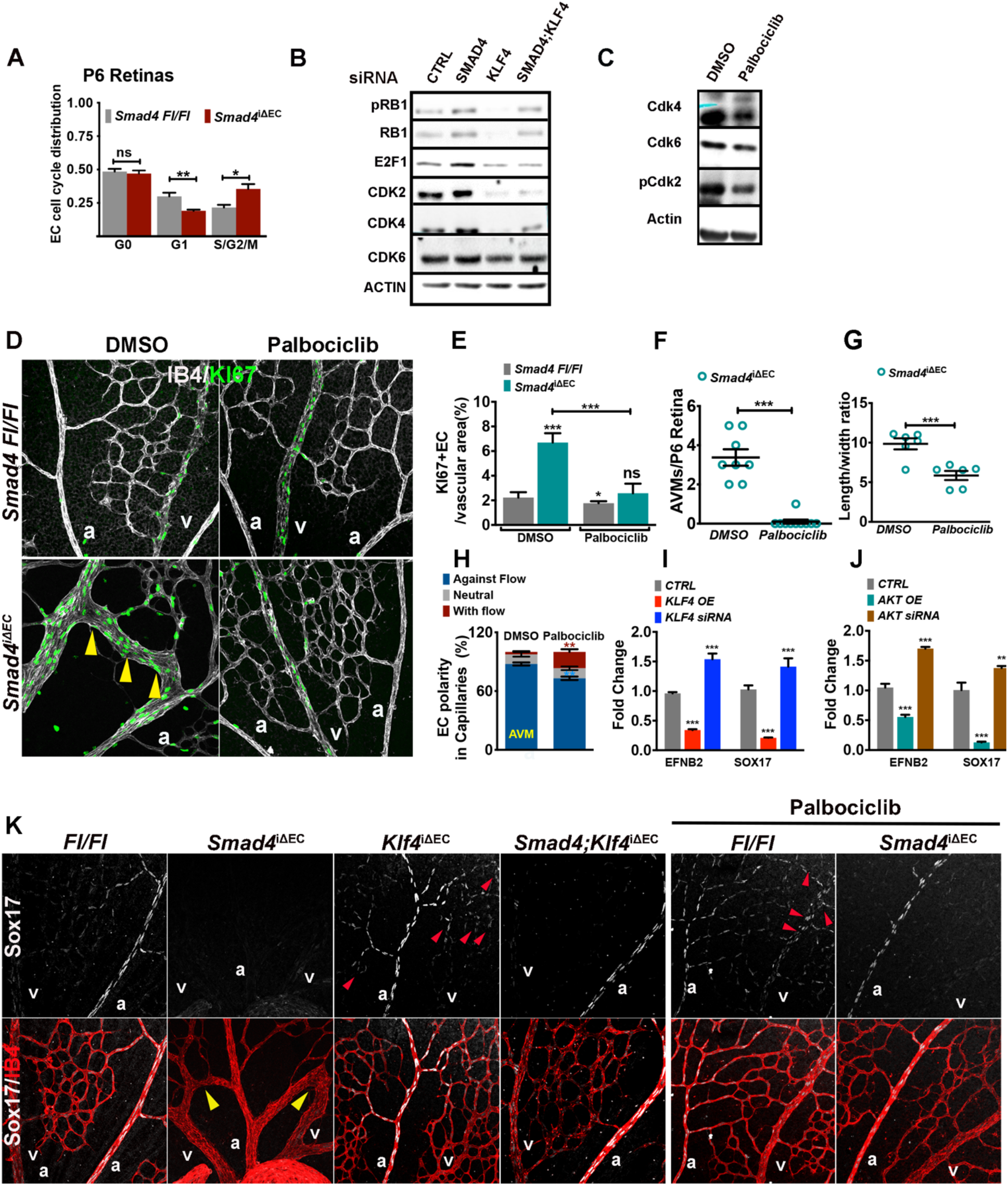
EC proliferation triggers AVM formation in *Smad4* deficient AVMs. (**A**) FACS analysis to assess cell cycle distribution in ECs from *Smad4 Fl/Fl and Smad4*^iΔEC^ P6 retinas. (**B**) WB for the indicated proteins of *CTRL, SMAD4, KLF4* and *SMAD4;KLF4* siRNAs HUVECs. (**C**) WB for the indicated proteins of whole lung lysates from pups treated with DMSO and Palbociclib. (**D**) Confocal images of P6 Smad4 *Fl/Fl* and *Smad4*^i∆EC^ retinas treated with DMSO or Palbociclib labeled for IB4 (white) and KI67 (green). Yellow arrows mark the KI67+ ECs within the AVMs. (**E**) Quantification of the number of KI67+ ECs per vascular area (%) in the vascular plexus. (**F**) Quantification of the number of AVMs in DMSO versus Palbociclib treated *Smad4*^iΔEC^ retinas. (**G**) Quantification of the ratio length/width in *Smad4*^i∆EC^ retinas DMSO or Palbociclib treated. (**H**) Quantification of the EC polarity in *Smad4*^iΔEC^ retinas DMSO or Palbociclib treated. (**I,J**) RT-PCR for *EPHRINB2* and *SOX17* in *CTRL, KLF4* siRNAs or KLF4*OE* and *CTRL, AKT* siRNAs and AKT*OE* HUVECs. (**K**) Confocal images of labeled retinas for Sox17 (white) and IB4 (red) from the indicated genotypes. Scale Bars: 100μm in **D,K**. Error bars: s.e.m., n.s-non-significant, \**P*<0.05, \*\**P*<0.01, \*\*\**P*<0.001, student T test. **a**: artery, **v**: vein.

Cell cycle progression is tightly regulated by members of the cyclins and cyclin dependent kinase (CDK) family. To identify dysregulated cell cycle regulators upon SMAD4 and KLF4 inactivation we performed WB analysis to multiple cell cycle regulators. *SMAD4*KD increased phosphorylation of Retinoblastoma (pRB1) and expression of E2F1 transcription factor, crucial regulators of cell cycle progression into S phase^26^ together with elevated CDK4, CDK6 and CDK2 protein levels (**Figure 7B**). Opposingly, *KLF4* inactivation alone led to reduced levels in all the main cell cycle regulators, suggesting cell cycle arrest. Interestingly, *KLF4*KD normalized the levels of pRB1, E2F1, CDK4 and CDK6 in *SMAD4*KD cells (**Figure 7B**).

To further test whether effects on cell cycle are the main drivers of AVMs, we treated *Smad4 Fl/Fl* and *Smad4*^iΔEC^ pups with Palbociclib, an inhibitor of CDK4/6 activity that efficiently blocks cell cycle progression^27^. We first confirmed efficacy of Palbociclib in lungs isolated from treated pups. In contrast to PBS treated pups, Palbociclib treatment decreased the expression of Cdk4 and Cdk6 and inhibited activation of Cdk2 (p-Cdk2) (**Figure 7C**).

In retinas labelled for IB4 and KI67, Palbociclib treatment decreased EC proliferation as quantified by the number of KI67+ ECs per vascular area in both *Control* and *Smad4*^iΔEC^ retinas (**Figure 7D**; quantified in **7E**) and significantly rescued the number of AVMs (**Figure 7D**; quantified in **7F**). We further assessed the impact of Palbociclib on morphological responses in *Smad4*ECKo retinas. Interestingly, similarly to KLF4-PI3K inhibition, Palbociclib treatment efficiently normalized the length/width ratio of *Smad4*ECko (**Figure 7G**) and also blunted the elevated orientation against the flow direction (**Figure 7H**). Taken together, these results suggest that increased cell proliferation as a result of flow-induced excessive KLF4-AKT-CDKs drives AVM formation.

Cell cycle arrest associated with physiological high flow is a prerequisite for maintaining arterial identity^27^. Conversely, dysregulated BMP9-SMAD4 signaling leads to loss of arterial and gain of venous identity^9,14,28^. Palbociclib-induced cell cycle arrest was reported to induce arterial markers^27^. To further untangle the connection between EC hyperproliferation and arterial-venous specification in this context, we analyzed HUVECs depleted or overexpressing *KLF4* or *AKT* for changes in arterial identity. RT-PCR identified a significant increase in expression of the arterial markers *EPHNB2* and *SOX17* upon knockdown of KLF4 and AKT, and decreases upon *KLF4* or *AKT OE* (**Figure 7I,J**).

To test these results *in vivo*, we labeled retinas for the arterial marker Sox17 (**Figure 7K**). In *Ctrl* retinas, Sox17 was confined to ECs in main arteries and a few arterioles. In AVMs in *Smad4-*deficient mice, Sox17 expression was completely abrogated. In *Klf4*ECko retinas, Sox17 expression expanded towards the vein and capillary ECs. *Klf4* inactivation in *Smad4*^iΔEC^ retinas largely rescued *Sox17* expression in arteries. Palbociclib treatment led to even greater expansion of Sox17 expression into capillary and venous ECs (**Figure 7K**). Collectively, these results suggest that increased EC proliferation-mediated loss of arterial identity is a central cell event in AVM formation.

## Discussion

It has been proposed that blood flow is ‘a second hit’ in HHT, as murine AVMs develop in regions of high shear stress^6,9^, but the mechanisms by which shear stress contributes to AVM pathogenesis remained largely undefined. ECs display an intrinsic set-point for physiological flow-induced shear stress that determines the signaling and gene expression outputs that control EC phenotype. VEGFR3 expression levels is one factor that can determine shear stress set-point for different types of vessels^5^. Also, non-canonical WNT signaling was proposed to modulate axial polarity set-point to control vessel regression in low flow regions^29^. We report here that loss of Smad4 in mouse ECs increased sensitivity to FSS with enhanced elongation and polarization in FSS together with diminished FSS-mediated cell cycle blockade and loss of EC arterial fate. Thus, Smad4 signaling is a novel mechanism that “sets the set-point” for high flow-mediated EC quiescence responses, e.g elongation, alignment and orientation. Smad4 is also critical for FSS-mediated growth suppression and arterial EC fate, though it remains to be determined if these events are also linked to disruption in the set point.

We previously reported that loss of BMP9-SMAD4 signaling potentiates PI3K/AKT activation^9,14^. Herein, we provide genetic evidence that PI3K/Akt activation is downstream of excessive Klf4 induction in *Smad4 ECko* or *KD* ECs, and that both Klf4 and PI3K/Akt are required for AVM formation. Interestingly, identification of excessive Klf4 and PI3K/Akt as a critical pathways in high-flow AVMs suggests mechanistic similarities with low-flow cerebral cavernous malformations (CCMs) and venous malformations (VM). In CCM lesions, malformations are initiated by mutation of *CCM1,2* or *3*, but GOF mutations in PI3K act as a third genetic hit (after clonal loss of the 2^nd^ CCM allele)^30^. Klf4 and its close homolog Klf2 are also highly induced in CCMs and contribute to lesion formation^31^. The low flow-VM are also a result of increased Akt function due to LOF mutations in PI3K, PTEN or TEK genes^32^.

It is generally assumed that the mechanisms of AVM formation are similar if not identical, in HHT1, HHT2 and JP-HHT. Our data, however, show that effects on EC polarization are essentially opposite. On one hand, *Smad4* LOF increased elongation and alignment in response to FSS, and polarization against the flow. By contrast, in *Eng* and *Alk1* LOF mice, ECs failure to polarize and migrate against the flow was proposed to promote AVM formation^8,15^. Similar cellular defects were suggested to be responsible for increased coronary arteries upon inactivation of embryonic *Smad4* in sinus venosus^33^. These correlations are intriguing, however, there is yet no functional or molecular evidence for its direct implication in AVM pathogenesis.

Searching for other mechanisms that may underly AVM development, deregulated proliferation and arterial-venous identity are attractive candidates, as lesions require both increased cell number and direct contact of arterial and venous ECs. Previous work showed loss of arterial identity and gain of venous markers in AVMs^9^, which contained exclusively venous-like ECs^4^. Additionally, arterial identity is linked to cell cycle arrest^34^. Thus, loss of cell cycle arrest due to loss of the BMP9/10-Alk1/Eng-Smad1/5 pathway could also lead to diminished expression of arterial identity genes that mediate repulsive interactions with venous ECs. These data do not rule out the possibility that loss of flow-driven migration is important for lesions driven by mutation of *Alk1* or *Eng*, implying distinct mechanisms for AVM formation due to different mutations. But at present, the hypothesis that migration direction is not the key event triggering AVMs appears both simpler and consistent with older studies demonstrating that depending on variables such as animal species, age and specific vascular location, ECs can migrate either against or with the flow^2,35^.

FSS induces cell cycle arrest in late G1 to enable maintenance of arterial identity via a Notch-Cx37-p27 signaling axis^27^. Notch and Smad1/5 co-regulate a number of genes, raising the possibilities that these two pathways function together^36^. We now report that Smad4 is also required for flow-induced cell cycle arrest-mediated arterial identity by restricting flow-induced Klf4-PI3K/Akt-CDK signaling. Genetic loss of Klf4 or pharmacological inhibition of PI3K or CDK4/6 rescues EC proliferation and restores arterial identity leading to EC normalization in *Smad4ECKO* retinas. Our findings support the concept that in Smad4 deficient EC, AVMs arise from loss of shear stress-mediated repression of EC proliferation and arterial identity due to excessive flow-induced Klf4-PI3K/Akt-Cdk (**Figure 8**).

**Figure 8.**
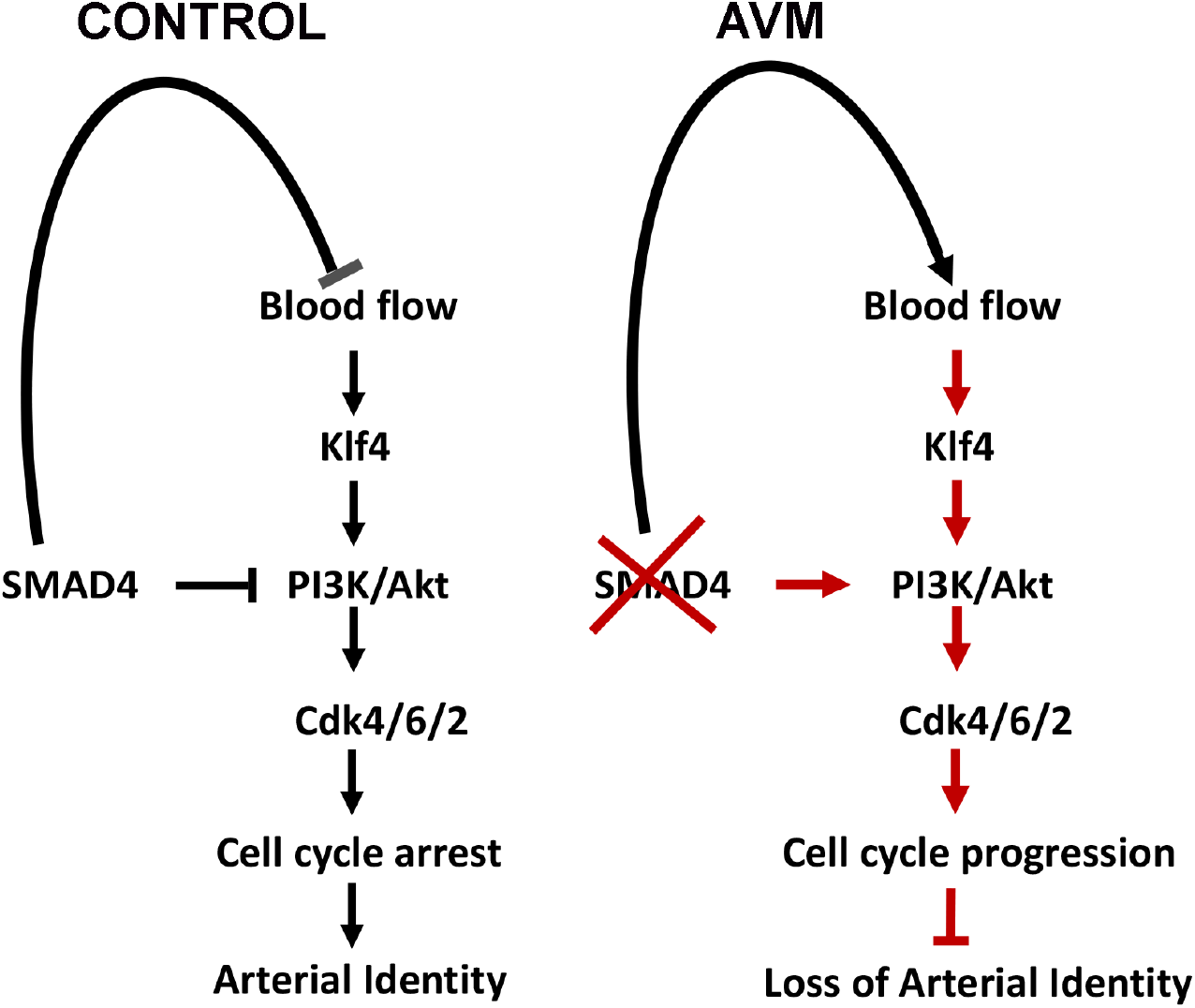
Working model for Smad4-Flow crosstalk in maintaining EC quiescence. SMAD4 restricts flow-induced activation of Klf4-Akt-Cdks to promote endothelial cell cycle arrest-mediated arterial identity. Loss of Smad4 leads to overactivation of flow-induced Klf4-Akt-Cdks leading to increased EC proliferation mediated loss of arterial identity and AVM formation.

These studies raise a number of new questions. Why does Klf2/4 induced by physiological flow stabilize vessels whereas higher levels promote cell proliferation and contribute to pathologies? Klf4 amplification of Akt activation is likely key but this mechanism is also unknown. Further work will be required to elucidate mechanism by which Smad4 “sets the set-point” for FSS-mediated EC responses to maintain EC quiescence and how Smad4 limits flow-induced activation of Klf4 in ECs.

Although AVMs in HHT patients form later in life, this is likely due to the requirement for second hit mutations that yield the homozygous mutant clones that initiate lesions. These pathways are thus likely to be relevant to human disease that afflicts mature vasculature. Targeting the Klf4-PI3K/Akt-Cdks axis may be a novel approach for developing new therapeutics for vascular malformations.

## Acknowledgements

The authors would like to thank Dr. Thomas Wielland (University Medical Center Mannheim) for sharing reagents. We would like to thank Stefanie Uhlig, Flow Core Mannheim and Institute of Transfusion Medicine and Immunology, for excellent support.

## References

1 Baeyens, N., Bandyopadhyay, C., Coon, B. G., Yun, S. & Schwartz, M. A. Endothelial fluid shear stress sensing in vascular health and disease. J Clin Invest 126, 821–828, doi:10.1172/JCI83083 (2016).

2 McCue, S. et al. Shear stress regulates forward and reverse planar cell polarity of vascular endothelium in vivo and in vitro. Circ Res 98, 939–946, doi:10.1161/01.RES.0000216595.15868.55 (2006).

3 Franco, C. A. et al. Correction: dynamic endothelial cell rearrangements drive developmental vessel regression. PLoS Biol 13, e1002163, doi:10.1371/journal.pbio.1002163 (2015).

4 Lee, H. W. et al. Role of Venous Endothelial Cells in Developmental and Pathologic Angiogenesis. Circulation 144, 1308–1322, doi:10.1161/CIRCULATIONAHA.121.054071 (2021).

5 Baeyens, N. et al. Vascular remodeling is governed by a VEGFR3-dependent fluid shear stress set point. Elife 4, doi:ARTN e0464510.7554/eLife.04645 (2015).

6 Baeyens, N. et al. Defective fluid shear stress mechanotransduction mediates hereditary hemorrhagic telangiectasia. J Cell Biol 214, 807–816, doi:10.1083/jcb.201603106 (2016).

7 Deng, H. et al. Activation of Smad2/3 signaling by low fluid shear stress mediates artery inward remodeling. Proc Natl Acad Sci U S A 118, doi:10.1073/pnas.2105339118 (2021).

8 Jin, Y. et al. Endoglin prevents vascular malformation by regulating flow-induced cell migration and specification through VEGFR2 signalling. Nat Cell Biol 19, 639-+, doi:10.1038/ncb3534 (2017).

9 Ola, R. et al. SMAD4 Prevents Flow Induced Arteriovenous Malformations by Inhibiting Casein Kinase 2. Circulation 138, 2379–2394, doi:10.1161/CIRCULATIONAHA.118.033842 (2018).

10 Johnson, D. W. et al. Mutations in the activin receptor-like kinase 1 gene in hereditary haemorrhagic telangiectasia type 2. Nat Genet 13, 189–195, doi:10.1038/ng0696-189 (1996).

11 McAllister, K. A. et al. Endoglin, a TGF-beta binding protein of endothelial cells, is the gene for hereditary haemorrhagic telangiectasia type 1. Nat Genet 8, 345–351, doi:10.1038/ng1294-345 (1994).

12 Teekakirikul, P. et al. Thoracic aortic disease in two patients with juvenile polyposis syndrome and SMAD4 mutations. Am J Med Genet A 161A, 185–191, doi:10.1002/ajmg.a.35659 (2013).

13 Crist, A. M. et al. Angiopoietin-2 Inhibition Rescues Arteriovenous Malformation in a Smad4 Hereditary Hemorrhagic Telangiectasia Mouse Model. Circulation 139, 2049–2063, doi:10.1161/CIRCULATIONAHA.118.036952 (2019).

14 Ola, R. et al. PI3 kinase inhibition improves vascular malformations in mouse models of hereditary haemorrhagic telangiectasia. Nat Commun 7, 13650, doi:10.1038/ncomms13650 (2016).

15 Park, H. et al. Defective Flow-Migration Coupling Causes Arteriovenous Malformations in Hereditary Hemorrhagic Telangiectasia. Circulation 144, 805–822, doi:10.1161/CIRCULATIONAHA.120.053047 (2021).

16 Iriarte, A. et al. PI3K (Phosphatidylinositol 3-Kinase) Activation and Endothelial Cell Proliferation in Patients with Hemorrhagic Hereditary Telangiectasia Type 1. Cells-Basel 8, doi:ARTN 97110.3390/cells8090971 (2019).

17 Dimmeler, S., Assmus, B., Hermann, C., Haendeler, J. & Zeiher, A. M. Fluid shear stress stimulates phosphorylation of Akt in human endothelial cells: involvement in suppression of apoptosis. Circ Res 83, 334–341, doi:10.1161/01.res.83.3.334 (1998).

18 Sugden, W. W. et al. Endoglin controls blood vessel diameter through endothelial cell shape changes in response to haemodynamic cues. Nat Cell Biol 19, 653-+, doi:10.1038/ncb3528 (2017).

19 Akimoto, S., Mitsumata, M., Sasaguri, T. & Yoshida, Y. Laminar shear stress inhibits vascular endothelial cell proliferation by inducing cyclin-dependent kinase inhibitor p21(Sdi1/Cip1/Waf1). Circ Res 86, 185–190, doi:10.1161/01.res.86.2.185 (2000).

20 Hamik, A. et al. Kruppel-like factor 4 regulates endothelial inflammation. J Biol Chem 282, 13769–13779, doi:10.1074/jbc.M700078200 (2007).

21 Bernabeu, M. O. et al. Computer simulations reveal complex distribution of haemodynamic forces in a mouse retina model of angiogenesis. J R Soc Interface 11, doi:ARTN 2014054310.1098/rsif.2014.0543 (2014).

22 Nagaoka, T. & Yoshida, A. Noninvasive evaluation of wall shear stress on retinal microcirculation in humans. Invest Ophth Vis Sci 47, 1113–1119, doi:10.1167/iovs.05-0218 (2006).

23 Peacock, H. M. et al. Impaired SMAD1/5 Mechanotransduction and Cx37 (Connexin37) Expression Enable Pathological Vessel Enlargement and Shunting. Arterioscl Throm Vas 40, E87–E104, doi:10.1161/Atvbaha.119.313122 (2020).

24 Coon, B. G. et al. Intramembrane binding of VE-cadherin to VEGFR2 and VEGFR3 assembles the endothelial mechanosensory complex. Journal of Cell Biology 208, 975–986, doi:10.1083/jcb.201408103 (2015).

25 Tzima, E. et al. A mechanosensory complex that mediates the endothelial cell response to fluid shear stress. Nature 437, 426–431, doi:10.1038/nature03952 (2005).

26 DeGregori, J., Kowalik, T. & Nevins, J. R. Cellular targets for activation by the E2F1 transcription factor include DNA synthesis- and G1/S-regulatory genes. Mol Cell Biol 15, 4215–4224, doi:10.1128/MCB.15.8.4215 (1995).

27 Fang, J. S. et al. Shear-induced Notch-Cx37-p27 axis arrests endothelial cell cycle to enable arterial specification (vol 8, 2017). Nature Communications 9, doi:ARTN 72010.1038/s41467-018-03076-4 (2018).

28 Mahmoud, M. et al. Pathogenesis of Arteriovenous Malformations in the Absence of Endoglin. Circulation Research 106, 1425–U1270, doi:10.1161/Circresaha.109.211037 (2010).

29 Franco, C. A. et al. Non-canonical Wnt signalling modulates the endothelial shear stress flow sensor in vascular remodelling. Elife 5, e07727, doi:10.7554/eLife.07727 (2016).

30 Ren, A. A. et al. PIK3CA and CCM mutations fuel cavernomas through a cancer-like mechanism. Nature 594, 271–276, doi:10.1038/s41586-021-03562-8 (2021).

31 Zhou, Z. N. et al. Cerebral cavernous malformations arise from endothelial gain of MEKK3-KLF2/4 signalling. Nature 532, 122-+, doi:10.1038/nature17178 (2016).

32 Queisser, A., Seront, E., Boon, L. M. & Vikkula, M. Genetic Basis and Therapies for Vascular Anomalies. Circ Res 129, 155–173, doi:10.1161/CIRCRESAHA.121.318145 (2021).

33 Poduri, A. et al. Endothelial cells respond to the direction of mechanical stimuli through SMAD signaling to regulate coronary artery size. Development 144, 3241–3252, doi:10.1242/dev.150904 (2017).

34 Chavkin, N. W. et al. Endothelial Cell Cycle State Determines Propensity for Arterial-Venous Fate. bioRxiv, 2020.2008.2012.246512, doi:10.1101/2020.08.12.246512 (2020).

35 Kiosses, W. B., McKee, N. H. & Kalnins, V. I. Evidence for the migration of rat aortic endothelial cells toward the heart. Arterioscler Thromb Vasc Biol 17, 2891–2896, doi:10.1161/01.atv.17.11.2891 (1997).

36 Larrivee, B. et al. ALK1 signaling inhibits angiogenesis by cooperating with the Notch pathway. Dev Cell 22, 489–500, doi:10.1016/j.devcel.2012.02.005 (2012).

